# PhyloScore: Improved covariation score for RNA alignments using phylogeny

**DOI:** 10.64898/2026.10.20.760377

**Authors:** Benjamin Lieser, Johannes Söding

## Abstract

**Motivation:** Structured RNAs can carry out complex biological functions similar to proteins, but our understanding of RNA structure and function lags far behind that of proteins. Covariation signals arising from RNA evolution can be recovered from sequence alignments and provide valuable insights into structure and function. However, such signals can also arise purely as a result of phylogenetic relatedness. We aimed to develop a method which can make use of the phylogenetic tree to distinguish better between real and false signal.

**Results:** PhyloScore predicts the manually curated secondary structure annotations in Rfam more accurately than the best currently used scores. On 48 Rfam families with experimentally determined structures, PhyloScore increases the mean average precision for recovering the annotated base pairs from 0.679 (R-scape’s APC-corrected G-test) to 0.714 under cross-validation, and improves 41 of the 48 families. Its p-values are discriminative across families, making them valuable for testing whether a non-coding RNA has a conserved structure.

**Availability:** PhyloScore is available as a pip package *“phylo_score”* under MIT Licence. The code repository is at https://github.com/soedinglab/phylo_score. The version described in this paper, including test data, is archived at https://doi.org/10.5281/zenodo.23401723

## 1 Introduction

RNA structure prediction remains a significant challenge, as shown by the results of RNA-Puzzles Round V (Bu et al., 2025), where no method could reliably predict all the structures and the most successful methods included human expert knowledge. Of the 4,178 RNA families currently cataloged in Rfam, only 158 have experimentally determined structures (Ontiveros-Palacios et al., 2025), compared to more than 12,000 Pfam protein families (Paysan-Lafosse et al., 2025) with at least one solved structure (Dana et al., 2019). This scarcity makes evolutionary signals a particularly valuable source of structural information. If an RNA molecule’s structure is essential to its function, evolutionary traces of that structure should be detectable in sequence alignments. For example, positions that form a conserved Watson-Crick base pair should show strong correlation with one another due to compensatory mutations. Such signals can aid RNA analysis and structure prediction.

The input of our tool is a multiple sequence alignment of homologous RNA sequences. It outputs for each pair of homologous positions a covariation score and the associated p-value for this score. R-scape (Rivas et al., 2017) does the same thing. It supports multiple scores, but it found the average product corrected G-test to be the most accurate. The G-test is a variant of mutual information with special handling of gaps.

However, the G-test has a limitation: covariation between two aligned sites can also arise from phylogenetic relationships rather than genuine co-evolution (Dutheil, 2012). If two mutations arise together in a single ancestor, they will tend to co-occur in all of its descendants. Without accounting for the underlying phylogenetic tree, this can produce a strong covariation signal even in the absence of compensatory mutations. Existing covariation scores correct for phylogeny only in limited ways, such as down-weighting similar sequences, while still treating sequences as independent observations. This can lead to false signals, as illustrated in Figure 1. R-scape goes one step further and computes empirical p-values by sampling null sequences with the same phylogenetic dependencies as the input MSA, thereby avoiding inflated p-values but failing to correct their co-evolution score for the dependence.

**Figure 1:**
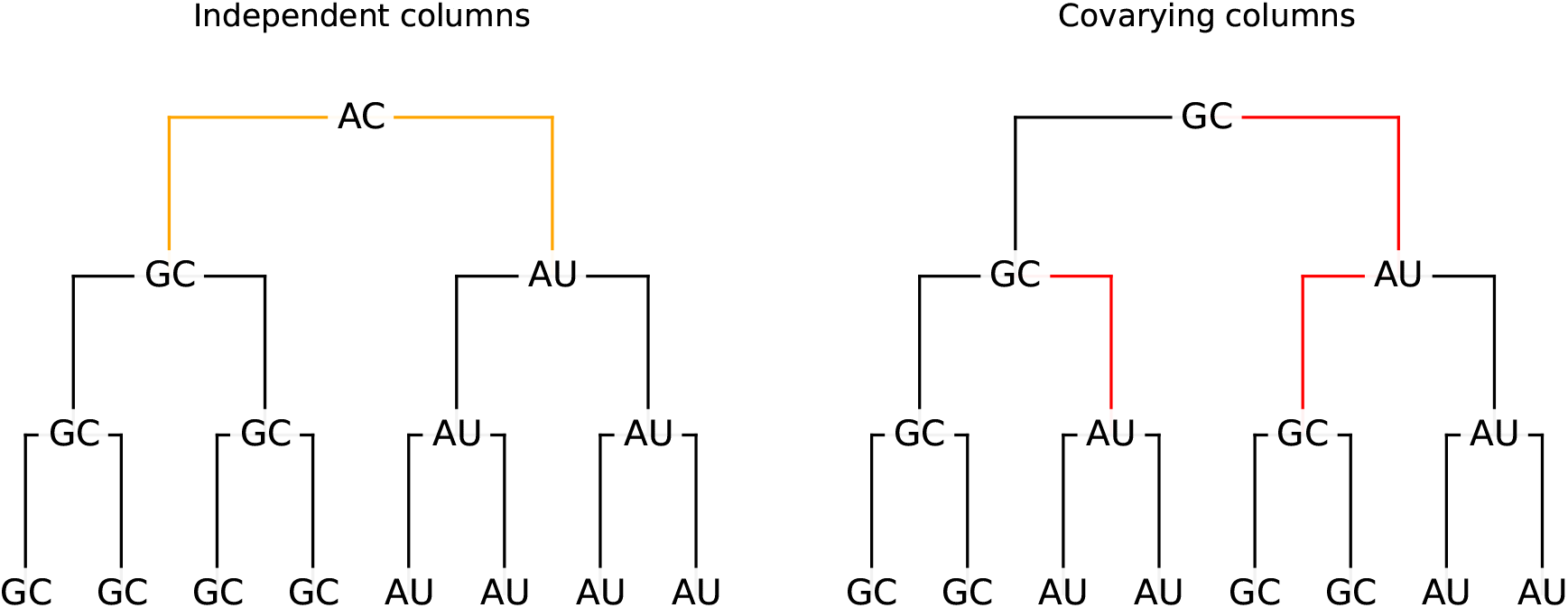
Independent evolution can produce strong spurious covariation signals. Left: Nucleotides at two independent positions at which two mutations occur near the root. In the multiple sequence alignment, the columns will be correlated despite the absence of any compensatory mutations. Right: compensatory mutations produce the identical set of sequences as on the left. Our goal is to distinguish between these two scenarios using phylogenetic information.

Rivas et al. (2017) excluded two phylogeny-aware methods (Akmaev et al., 2000; Yeang et al., 2007) from their benchmark, noting that neither has a publicly available implementation and both are computationally expensive. Related phylogenetic models of co-evolving sites have also been developed for protein and nucleotide sequences (Pollock et al., 1999; Dib et al., 2014). We aimed to make this approach using probabilistic phylogenetic models of co-evolution both accessible and computationally efficient. We show that incorporating phylogenetic information improves covariation scores while keeping computational cost manageable.

## 2 Method

We start from an RNA alignment and predict a phylogenetic tree from it using the GTR+G model (Tavaré, 1986; Yang, 1994) in RAxML-NG (Kozlov et al., 2019).

We specify parametrized substitution models on the 16 pairs of nucleotides (AA, AC, AG, AU, CA, …) and compare the resulting likelihood of a model with independent evolution against a model which models co-evolving pairs of positions.

Given a tree, a model and an alignment, the Felsenstein algorithm (Felsenstein, 1981) can efficiently calculate the likelihood of this alignment under the tree and the model. PhyloGrad (Lieser et al., 2025) can efficiently compute gradients of the log-likelihood with respect to the model parameters, which allows us to optimize the model parameters efficiently.

### 2.1 General time reversible model

The general time reversible model (GTR) (Tavaré, 1986) is the basis for our two models. A Continuous Time Markov Chain is described by its rate matrix *Q* such that the probability of changing from a pair of nucleotides *a* to a pair *b* in time *t* is given by exp(*tQ*)_*ab*_. To make these probabilities properly normalized, the rows of the rate matrix have to sum to 0 and only the diagonal elements are negative.

For time reversibility, we require *q*_*ab*_*π*_*a*_ = *q*_*ba*_*π*_*b*_, where *π* is the equilibrium distribution of the Markov chain. We achieve this by parameterizing *Q* as 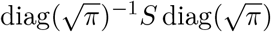, where *S* is a symmetric matrix with non-negative entries and 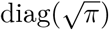 denotes the diagonal matrix with Entries 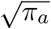. Diagonal elements of S are unused, because the diagonal elements of *Q* are already determined by the off-diagonal elements. We re-parametrize 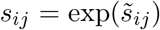 and 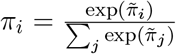 to enforce non-negativity and normalization. We do not normalize, so the model can fit itself to the given branch lengths.

### 2.2 Model for independently evolving positions

We use two independent 4-state (A,C,G,U) GTR models, which have parameters 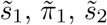 and 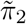. To make this model analogous to the co-evolution model, we merge the two independent models into the equivalent 16-state model. This is done via *π*_*ab*_ = *π*_1*a*_*π*_2*b*_ and

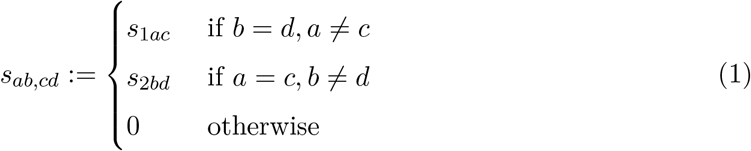

A proof of the equivalence to two independent chains can be found in the supplement. Note that the diagonal elements do not need to be specified. We subtract from the log-likelihood an *L*_2_ regularization term

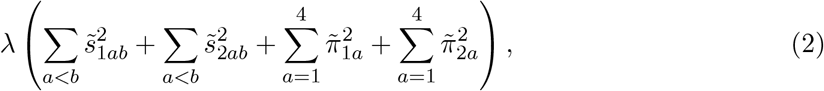

where *λ ≥* 0 determines the regularization strength.

### 2.3 Model for co-evolving positions

The *S* matrix has 16 *·* 15*/*2 = 120 parameters which are too many to reliably estimate. We therefore use the same *S* matrix as in the independent model eq. (1). The 16 probabilities of *π* can be learned freely, which allows the model to fit Watson-Crick pairing or other pairing preferences.

Parameters of the independent model also describe a valid co-evolution model. The 16 equilibrium parameters are just given by *π*_*ab*_ = *π*_1*a*_*π*_2*b*_. We want the regularization to match the one of the independent model in this case, so we calculate the eight marginal probabilities *π*_1_ *∈* ℝ^4^ and *π*_2_ *∈* ℝ^4^ from *π ∈* ℝ^16^ and use the same regularization terms as in equation (2). In addition, we add a mutual information penalty *λ*_MI_MI(*π*), with 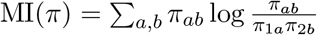.

### 2.4 The score

We use two different approaches to derive a score from these two models.

#### log-likelihood difference

This is the log-likelihood of the co-evolution model minus the log-likelihood of the independent model. It ignores the regularization terms, which are only used during the model fit. This can result in negative scores.

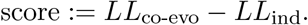

#### regularized log-likelihood difference

It uses:

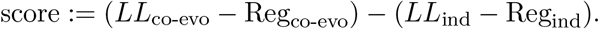

This is the difference of the two functions which are maximized, so the score will always be non-negative, because we initialize the optimization of the co-evolution model with the independent model and the independent model is contained in the co-evolution model.

### 2.5 Average product and sum correction

The average product correction (Dunn et al., 2008) is a common correction applied after estimation of co-evolution scores between pairs of an alignment. The log-likelihood difference score can be negative, so if that happens we subtract the smallest score from all scores before applying the average product correction. Let *C*_*ij*_ = *C*_*ji*_ be the score between columns *i* and *j* and *L* be the number of columns.

We define 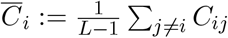 and 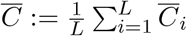,

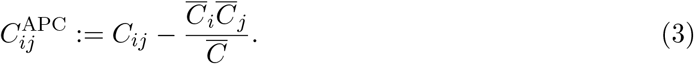

Another possible correction is the average sum correction given by

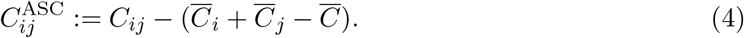

### 2.6 Gap Treatment

If either of two columns has a gap in a given sequence, we remove that sequence from the likelihood calculation.

### 2.7 P-Values

We calculate two different p-values. The fast p-values use the results from the original alignment. If we have *L* columns in the alignment, we have *L*(*L −* 1)*/*2 many pairs. At most *L/*2 of the pairs can be paired. So if we remove the highest *L/*2 scores we should have only non-paired pairs left which are used to train the null model.

The null p-values come from fitting the null model on scores from an R-scape null alignment. This works slightly better, but doubles the runtime because scores also need to be calculated for the null alignment. In both cases we fit a gamma distribution with location parameter using `scipy.stats.gamma.fit` (Virtanen et al., 2020) as a null model.

### 2.8 Implementation

We implemented the model fitting part in Rust. It uses Adam (Kingma and Ba, 2015) for optimization and computes the gradients using PhyloGrad (Lieser et al., 2025), which we have integrated into the automatic differentiation framework `candle` (Hugging Face, 2023). PhyloScore comes as an MIT-licensed Python package, which can be installed with `pip` (`pip install phylo_score` and then run with `python -m phylo_score`). For ease of use we re-implemented R-scape’s null alignment generation.

## 3 Results

### 3.1 Data

We test on seed alignments of the Rfam database (Ontiveros-Palacios et al., 2025) (version 15.1). These seed alignments are manually curated and have a consensus secondary structure annotation. This consensus annotation is the result of co-evolution scores, experimental structures, predicted structures and human expert analysis. For our benchmark, we only consider families with available experimental structure, which should result in more accurate annotations. We consider only seed-alignments with at least 100 sequences. Both conditions leave us with 49 families. We excluded RF02541 (Bacterial large subunit ribosomal RNA) because of its size of 2925 nucleotides.

We cannot say for sure whether the secondary structure annotations are accurate. Also, co-evolution can be driven by factors other than secondary structure. Despite this, we think it is a good benchmark to evaluate co-evolution scores because the annotated base pairs will have a significantly higher chance of having co-evolved.

### 3.2 Benchmark Method

For each seed alignment, we take all possible pairs of columns and classify them according to whether they have a pairing in the consensus annotation (label 1) or not (label 0). Columns with a gap in the consensus annotation are removed from the analysis. These are typically highly gapped columns in the alignment. We then compute the precision-recall curve and compute the area under this curve, often called the average precision. Because of the extremely unbalanced dataset (the vast majority of pairs will be unpaired), the ROC curve is unsuitable but precision-recall is robust against this. For the G-test and other already used scores we use R-scape (Rivas et al., 2017) (version 2.6.4) to compute them. We use version 1.5.5 of CoMap (Dutheil et al., 2005).

We did not include Coev (Dib et al., 2014), which also compares an independent and a co-evolution substitution model on the 16 nucleotide pairs. Its public implementation appears to be no longer maintained and crashes with a segmentation fault on its own example data. We also did not include the maximum likelihood method of Pollock et al. (1999) because it has no public source code available.

### 3.3 CoMap Results

CoMap could not complete on RF00005 and RF03064 because of a likelihood underflow. This makes it not directly comparable to the R-scape and PhyloScore.

Doing a comparison of the mean average precision over the 46 families it runs on gives us the following values:

**Table 1:** Mean average precision over the 46 families that CoMap could process. OMES is the worst performing R-scape score except for CCF (which is available in the code but not described in the paper)

| Method | Mean average precision |
| --- | --- |
| PhyloScore | 0.714 |
| R-scape GTp | 0.667 |
| R-scape GT | 0.602 |
| R-scape OMES | 0.493 |
| CoMap (with APC) | 0.374 |
| CoMap p-value | 0.319 |
| CoMap | 0.125 |

Because it performs so much worse than R-scape on our data, we will benchmark PhyloScore only against R-scape.

### 3.4 Cross-Validation

Our method has hyperparameters to be fit, so to make sure we have a fair benchmark we use cross-validation. These hyperparameters are the regularization strengths *λ* and *λ*_MI_, the type of score and the type of correction. We performed a grid search over the hyperparameters. The exact values used can be found in Table S1 and S2.

We split the families into five folds. For each hold-out fold we find the best hyperparameters on the four training folds and evaluate on the hold-out set. The metric is the mean average precision over the families. We can do the calculations for all hyperparameter combinations and families once, which allows us to do simulated cross-validation runs very quickly.

We do the same with R-scape using all the covariation scores available in R-scape. In our 1000 simulated cross-validations, only the GTp (the average product corrected G-Test) score was selected.

The histogram of 1000 five-fold cross-validation runs is shown in Figure 2. R-scape reaches a mean average precision of 0.679 in every run, while PhyloScore reaches between 0.710 and 0.715 (median 0.714). PhyloScore is therefore robustly better than R-scape, and the improvement is not due to fitting the hyperparameters on the benchmark families. Across all runs, the selected hyperparameters always used the regularized log-likelihood difference with *λ ∈ {*0.5, 1.0*}* and *λ*_MI_ *≤* 0.1.

**Figure 2:**
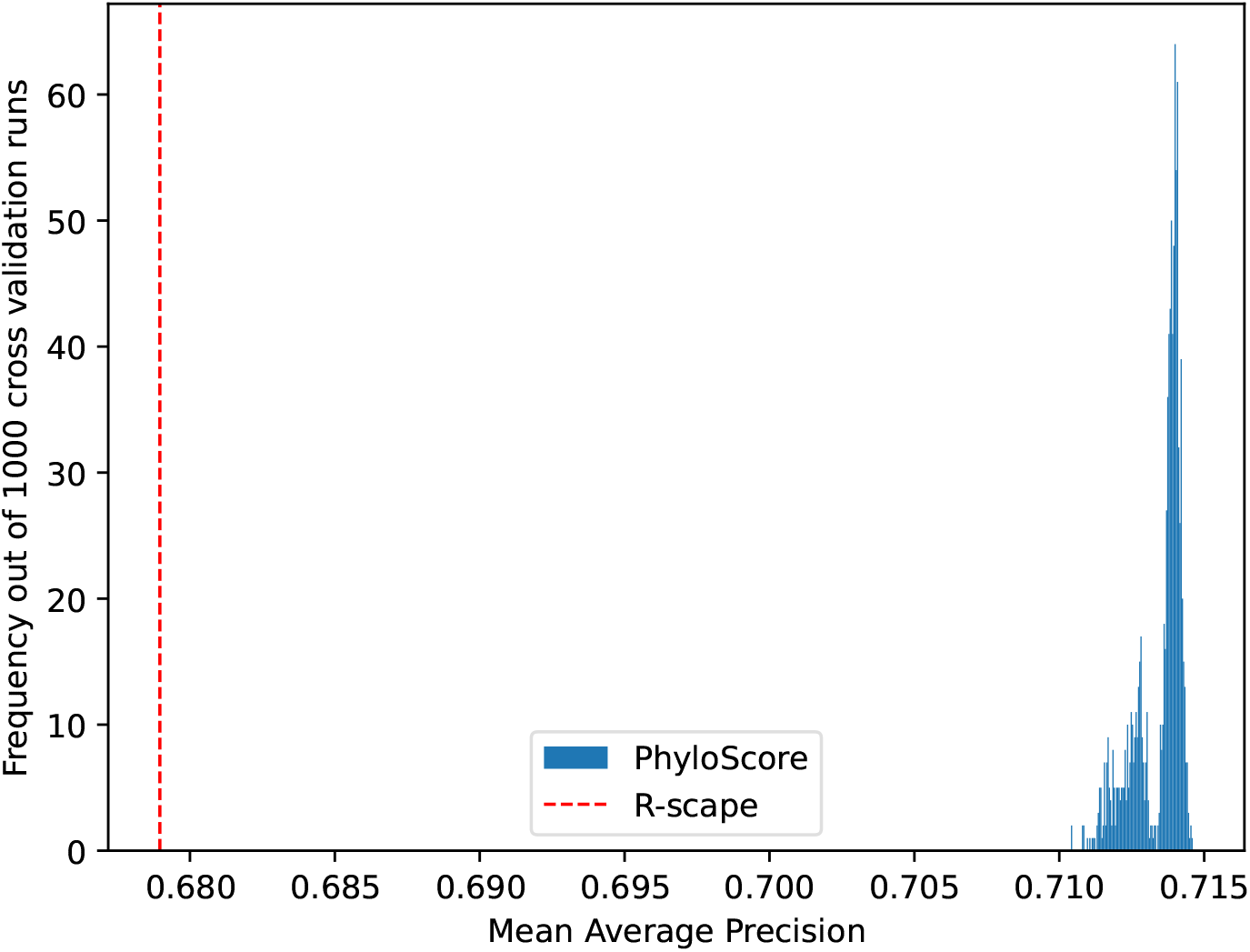
Mean of the average precision over all the families: The red line indicates R-scape’s fixed average precision in the cross-validation. The histogram shows the distribution of PhyloScore over 1000 five-fold cross-validations.

Figure 3 shows the average precision of R-scape and PhyloScore for the 48 families. PhyloScore improves the median cross-validated average precision for 41 families, by up to 0.148, and performs worse for 6 families; one family (RF00026) is predicted perfectly by both methods. Five of the six losses are below 0.03. The largest loss (RF00386, 0.251 versus 0.193) occurs in a family with low average precision for both methods, where covariation is not very predictive of the secondary structure in the first place.

**Figure 3:**
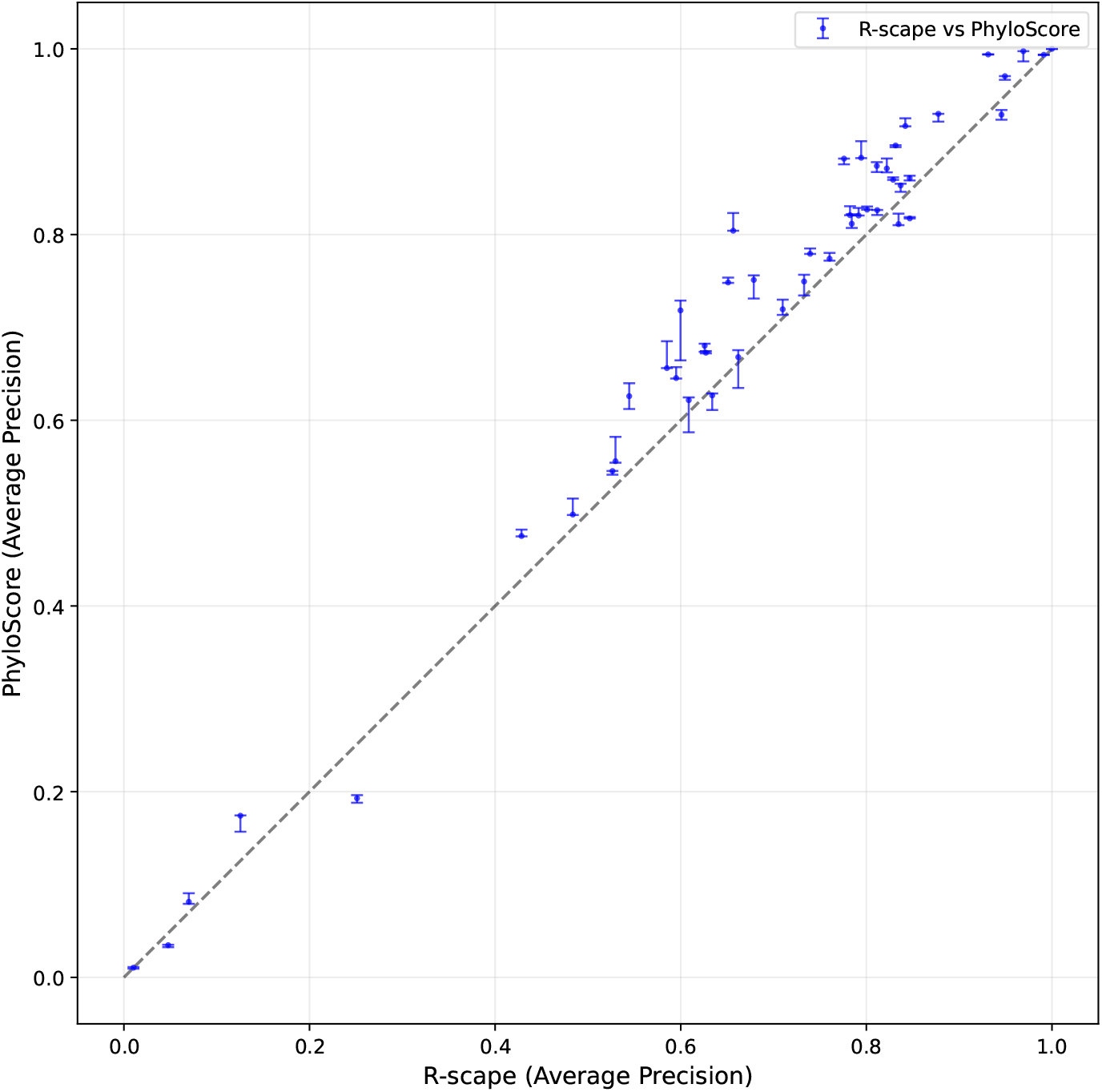
Average precision of PhyloScore and R-scape per family. The points are at the median of the cross validated average precisions. The error bars show the minimum and the maximum. For R-scape there is no variance in the cross-validation.

### 3.5 P-Values

In order to benchmark if the reported p-values are able to discriminate between pairs in different alignments, we combine all the families and see how well the p-values predict Rfam annotations when all the pairs of all the families are merged. If the scores were only comparable inside one alignment, then this would be detected in this benchmark.

For PhyloScore we use the best-performing hyperparameters from the cross-validation (*λ* = 0.5, *λ*_MI_ = 0.01, regularized likelihood difference, and average product correction).

The results are shown in Figure 4. PhyloScore is especially good in the low and high recall range compared to R-scape. In the medium recall range they have the same precision. Furthermore, the G-Test alone without the p-value adjustments from R-scape performs very poorly.

**Figure 4:**
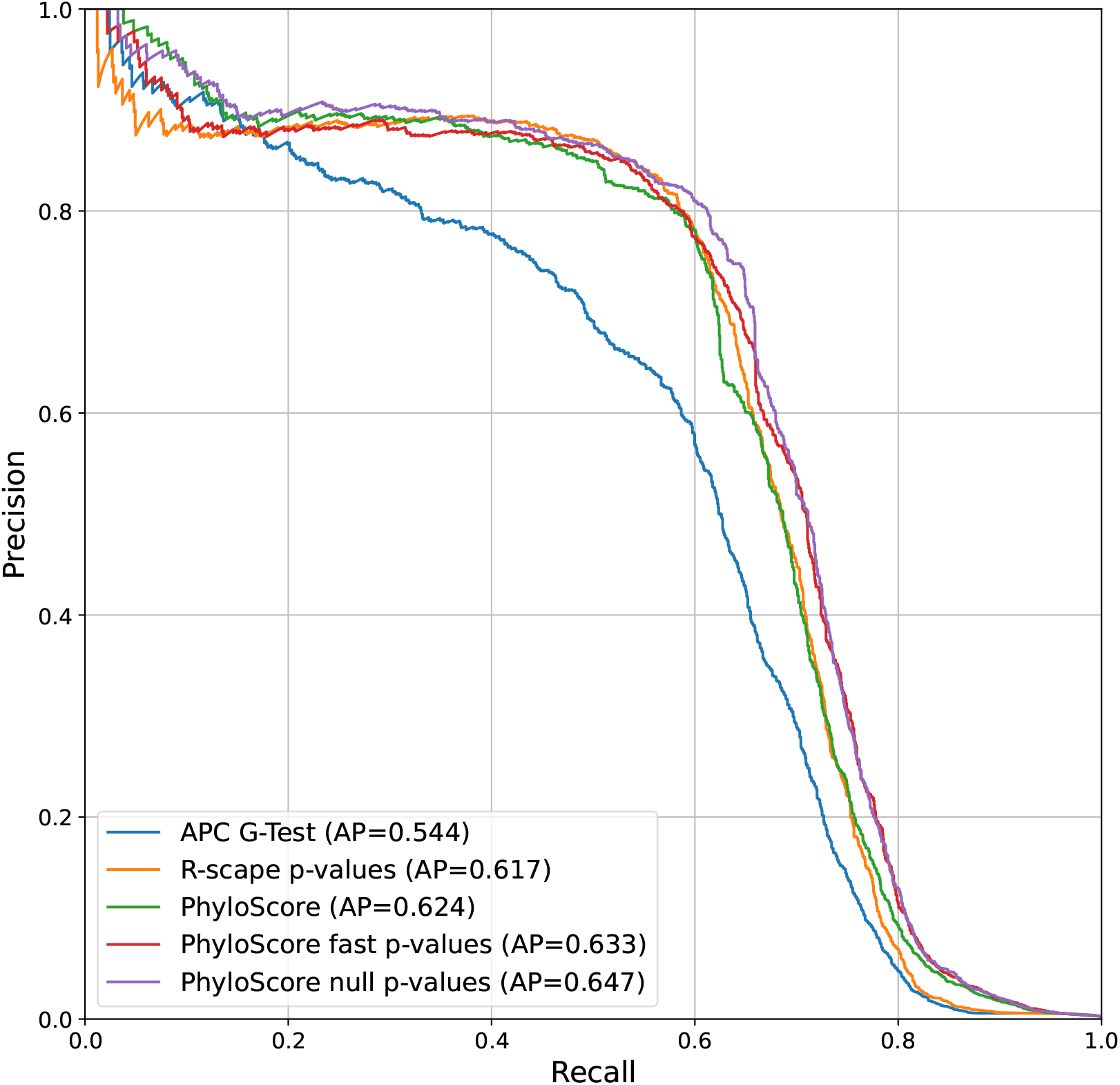
Precision Recall curve of all families combined for a) The average product corrected G-Test scores, b) The p-values calculated by R-scape of the G-Test, c) The PhyloScore score, d) The fast p-values from PhyloScore, e) The p-values on the null alignment from PhyloScore.

Note that the pure scores perform better in the low-recall range than the p-values for both R-scape and PhyloScore. This is probably because high absolute scores are meaningful and will be on the tail of the null distributions whereas small changes in the distribution parameters can have a big effect on the p-value.

## 4 Discussion

PhyloScore can detect covariation more sensitively than existing methods with a moderate computational cost. The explicit treatment of the phylogeny allows it to distinguish between correlation introduced by the phylogeny and correlation due to real co-evolution. Because its p-values are discriminative across alignments, it can be used as a more sensitive alternative to R-scape when testing for conserved secondary structure.

We use the annotations and alignments from Rfam, which are built with current scores already in use. So they might be biased towards these scores. Therefore, we probably overestimate R-scape performance here, which means the improvement of our method might be bigger than the benchmarking suggests.

R-scape is very fast to run, mostly less than 1 minute for all the tested families. PhyloScore is slower, for example, the 208-sequence alignment of RF00004 with its 18,336 pairs takes 72 minutes using 16 threads (4 pairs per second). We expect that this is not a big bottleneck for most applications where accuracy is more important than speed because there is only limited data available.

The G-Test needs the average product correction to work well. PhyloScore also profits from these corrections but significantly less. We can see these differences in Figures S2 and S3. It seems the average product correction can account for some false signal from phylogeny but not all.

## Conflicts of interest

The authors declare that they have no competing interests.

## Funding

This work was funded by the Max-Planck society.

## Data availability

The source code and test data underlying this article are available at Zenodo (https://doi.org/10.5281/zenodo.23401723) and GitHub (https://github.com/soedinglab/phylo_score).

## Author contributions statement

B.L. developed and implemented the model, designed and implemented the benchmarks and drafted the manuscript. J.S. supervised the work and reviewed the manuscript.

## Acknowledgments

B.L. thanks the International Max-Planck Research School for Genome Science for the support during his PhD. The AI assistant Claude (Anthropic) was used to reimplement R-scape’s null alignment generation in PhyloScore and write most of the CoMap benchmark code. It also helped edit the manuscript and check the references. The authors reviewed all AI-assisted code and text and take full responsibility for the content.

## Supplementary Material

### S1 Combining Models Proof

For the non-diagonal entries of the rate matrix *Q* we already have

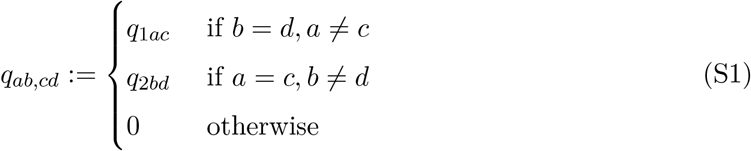

in case of two independent Markov models. See, for example, Yeang et al. (2007).

By definition we have 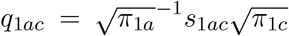 and 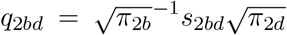 and 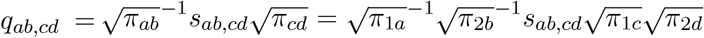. So for the case *b* = *d, a ≠ c* we get 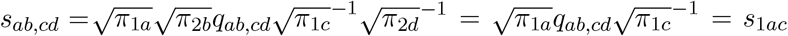. It is analogous for the other case, and it is trivial for the third case.

### S2 Hyperparameters used

**Table S1:** Hyperparameters for Cross-Validation of PhyloScore.

| Parameter | Values |
| --- | --- |
| $\lambda$ | {0.05, 0.1, 0.25, 0.5, 1.0, 2.5, 5.0} |
| $\lambda_{\text{MI}}$ | {0.0, 0.01, 0.1, 1.0, 10.0} |
| Score | {log-likelihood difference, regularized log-likelihood difference} |
| Correction | {average product, average sum, none} |

**Table S2:**
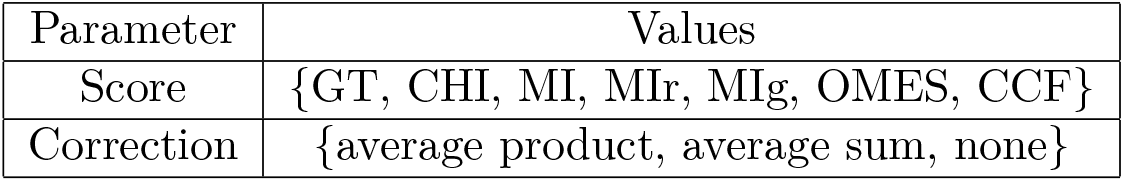
Hyperparameters for Cross-Validation of R-scape.

We could not get RAF and RAFS to run for R-scape.

### S3 Effect of the hyperparameters

From figure S1 we can see that the biggest effect comes from the regularization parameter *λ*. This is not surprising, because it controls how much the model parameter can fit the alignment. The mutual information penalty parameter *λ*_MI_ is less important.

The difference of the regularized likelihood was slightly better than just the difference in log likelihood.

There is almost no difference between the average product correction and the average sum correction. Both perform better than no correction, but only slightly.

**Figure S1:**
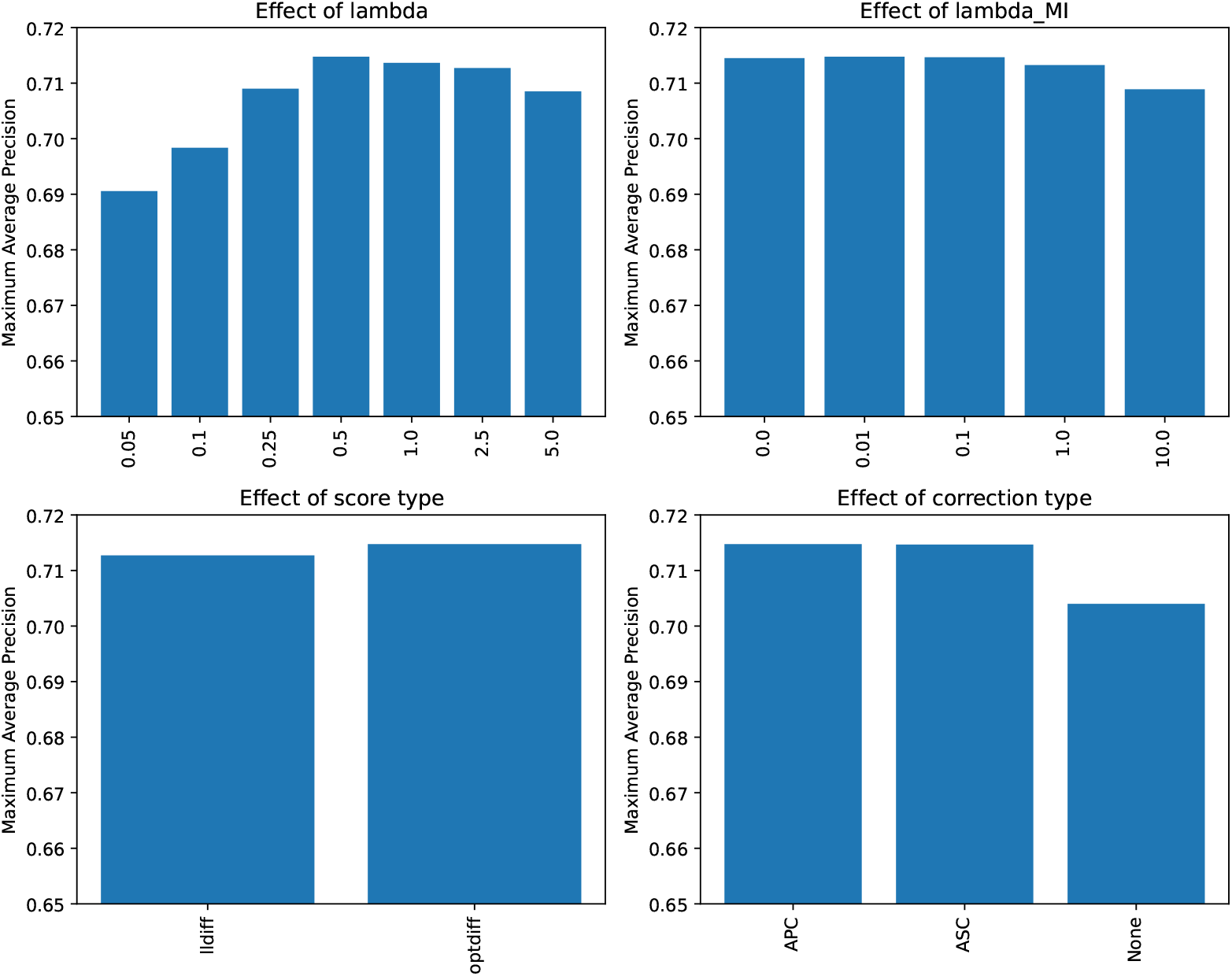
The plot shows the mean average precision over all families when fixing one hyper-parameter and maximizing over all others.

In comparison the corrected G-Test (0.679 average precision) performs significantly better than the uncorrected G-Test (0.613 average precision).

### S4 Effects of Corrections

PhyloScore profits significantly less from an average product/sum correction than the scores used in R-scape. These corrections might be able to remove a big part of the false signal induced from the phylogeny.

We can see this in Figures S2 and S3.

**Figure S2:**
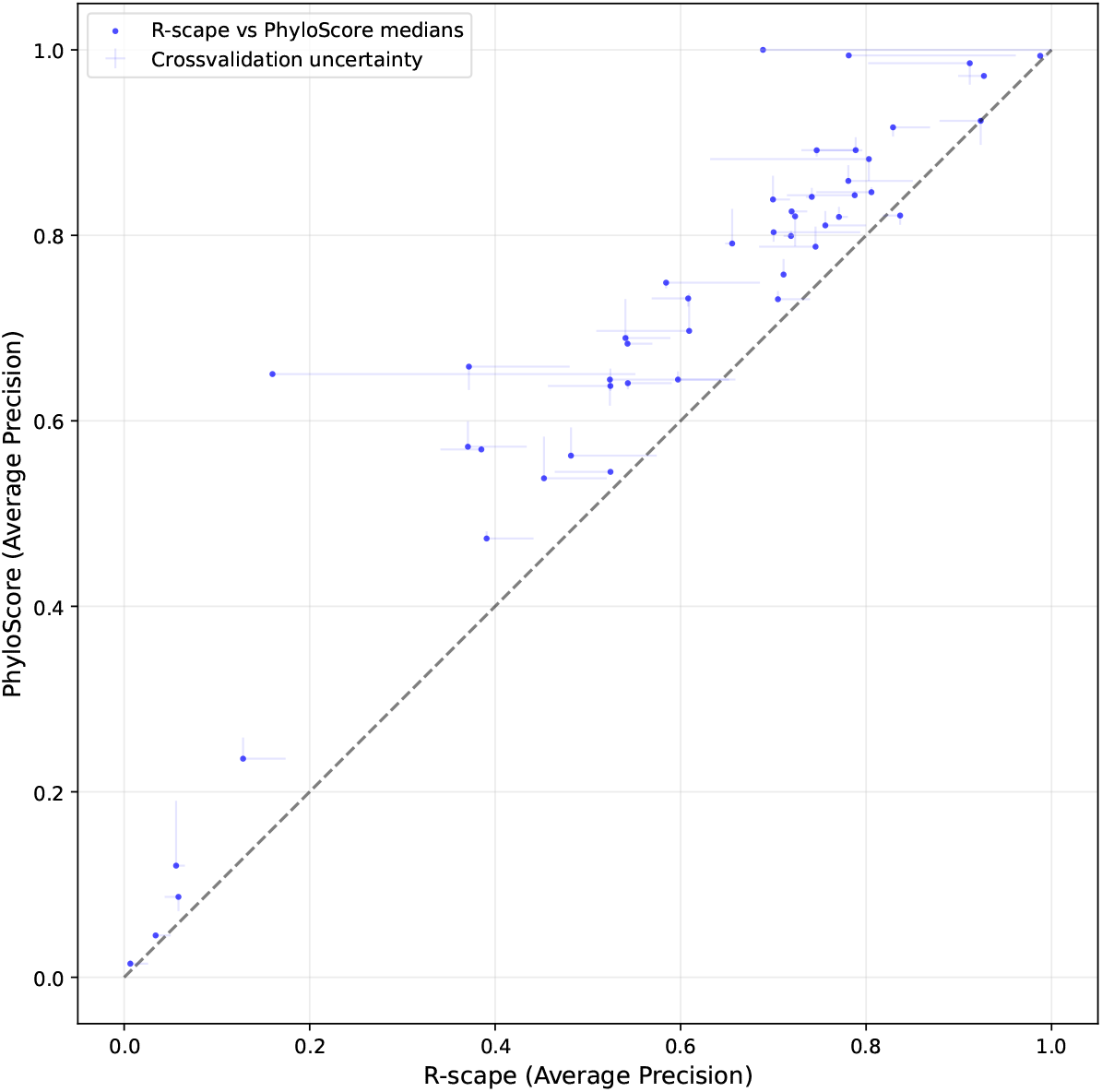
Average precision per family without corrections. The dots are the median results of the 1000 cross-validation runs, the bars give the best and worst result.

**Figure S3:**
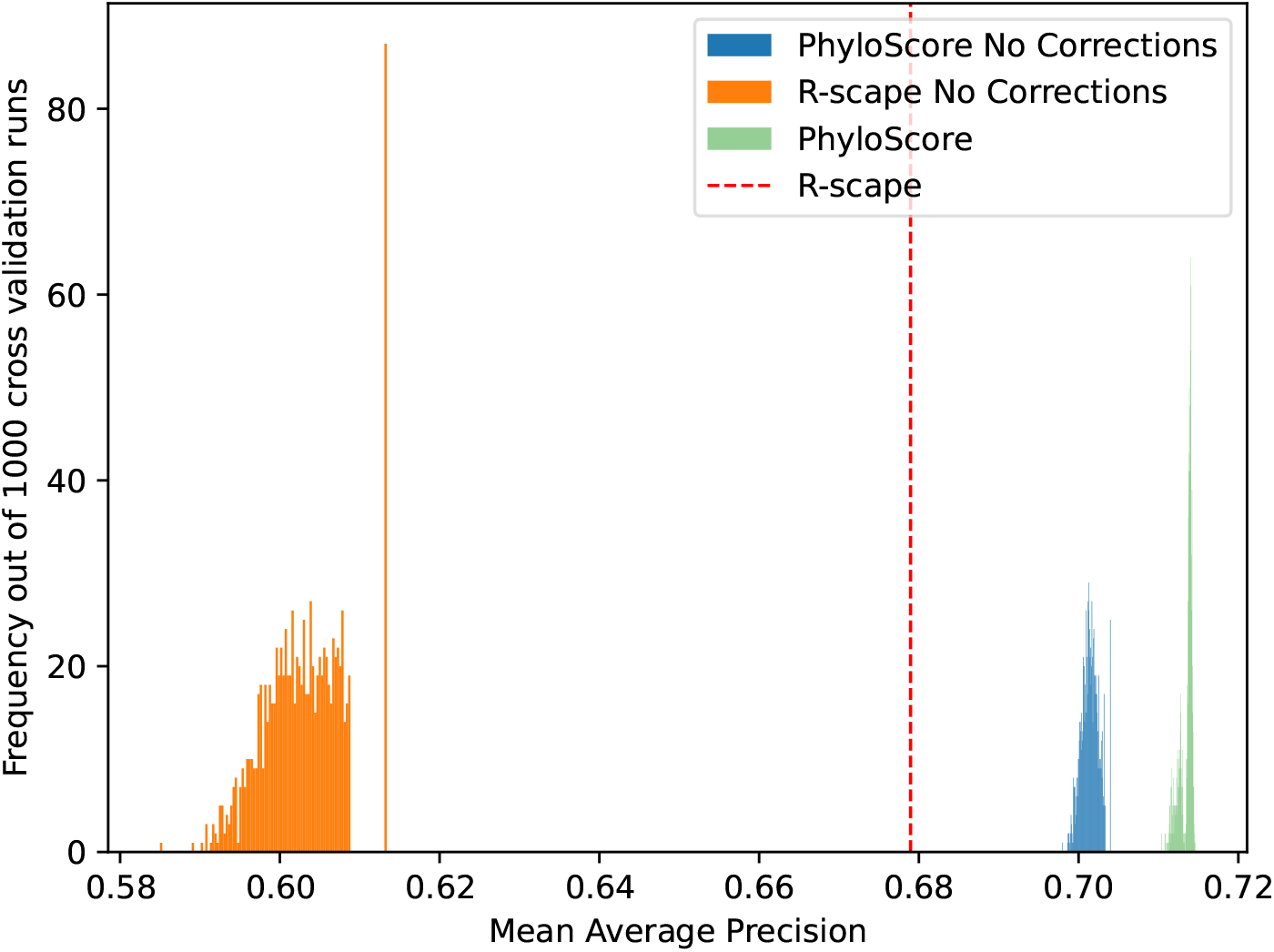
Overall average precision over 1000 simulated cross-validations without corrections. For R-scape the cross-validation now chooses either the G-Test or the Chi2 statistic, so there is some variance now.

## Notes

### Competing Interest Statement

The authors have declared no competing interest.

## References

V. R. Akmaev, S. T. Kelley, and G. D. Stormo. Phylogenetically enhanced statistical tools for RNA structure prediction. Bioinformatics, 16(6):501–512, 2000.

F. Bu, Y. Adam, R. W. Adamiak, M. Antczak, B. R. H. de Aquino, N. G. Badepally, R. T. Batey, E. F. Baulin, P. Boinski, M. J. Boniecki, et al. RNA-Puzzles Round V: blind predictions of 23 RNA structures. Nature Methods, 22(2):399–411, 2025.

J. M. Dana, A. Gutmanas, N. Tyagi, G. Qi, C. O’Donovan, M. Martin, and S. Velankar. SIFTS: updated structure integration with function, taxonomy and sequences resource allows 40-fold increase in coverage of structure-based annotations for proteins. Nucleic Acids Research, 47 (D1):D482–D489, 2019. doi: 10.1093/nar/gky1114.

L. Dib, D. Silvestro, and N. Salamin. Evolutionary footprint of coevolving positions in genes. Bioinformatics, 30(9):1241–1249, 2014. doi: 10.1093/bioinformatics/btu012.

S. D. Dunn, L. M. Wahl, and G. B. Gloor. Mutual information without the influence of phylogeny or entropy dramatically improves residue contact prediction. Bioinformatics, 24(3):333–340, 2008. doi: 10.1093/bioinformatics/btm604.

J. Dutheil, T. Pupko, A. Jean-Marie, and N. Galtier. A model-based approach for detecting coevolving positions in a molecule. Molecular Biology and Evolution, 22(9):1919–1928, 2005. doi: 10.1093/molbev/msi183.

J. Y. Dutheil. Detecting coevolving positions in a molecule: why and how to account for phylogeny. Briefings in Bioinformatics, 13(2):228–243, 2012. doi: 10.1093/bib/bbr048.

J. Felsenstein. Evolutionary trees from DNA sequences: a maximum likelihood approach. Journal of Molecular Evolution, 17(6):368–376, 1981. doi: 10.1007/BF01734359.

Hugging Face. Candle: a minimalist ML framework for Rust. https://github.com/huggingface/candle, 2023.

D. P. Kingma and J. Ba. Adam: A method for stochastic optimization. In 3rd International Conference on Learning Representations (ICLR), 2015. arXiv:1412.6980.

A. M. Kozlov, D. Darriba, T. Flouri, B. Morel, and A. Stamatakis. RAxML-NG: a fast, scalable and user-friendly tool for maximum likelihood phylogenetic inference. Bioinformatics, 35(21): 4453–4455, 2019. doi: 10.1093/bioinformatics/btz305.

B. Lieser, G. Belousov, and J. Söding. PhyloGrad: fast column-specific calculation of substitution model gradients. BMC Bioinformatics, 27:20, 2025. doi: 10.1186/s12859-025-06353-4.

N. Ontiveros-Palacios, E. Cooke, E. P. Nawrocki, S. Triebel, M. Marz, E. Rivas, S. Griffiths-Jones, A. I. Petrov, A. Bateman, and B. Sweeney. Rfam 15: RNA families database in 2025. Nucleic Acids Research, 53(D1):D258–D267, 2025.

T. Paysan-Lafosse, A. Andreeva, M. Blum, S. R. Chuguransky, T. Grego, B. L. Pinto, et al. The Pfam protein families database: embracing AI/ML. Nucleic Acids Research, 53(D1): D523–D534, 2025. doi: 10.1093/nar/gkae997.

D. D. Pollock, W. R. Taylor, and N. Goldman. Coevolving protein residues: maximum likelihood identification and relationship to structure. Journal of Molecular Biology, 287(1):187–198, 1999. doi: 10.1006/jmbi.1998.2601.

E. Rivas, J. Clements, and S. R. Eddy. A statistical test for conserved RNA structure shows lack of evidence for structure in lncrnas. Nature Methods, 14(1):45–48, 2017.

S. Tavaré. Some probabilistic and statistical problems in the analysis of DNA sequences. In Some Mathematical Questions in Biology: DNA Sequence Analysis, volume 17 of Lectures on Mathematics in the Life Sciences, pages 57–86. American Mathematical Society, 1986.

P. Virtanen, R. Gommers, T. E. Oliphant, et al. SciPy 1.0: fundamental algorithms for scientific computing in Python. Nature Methods, 17(3):261–272, 2020. doi: 10.1038/s41592-019-0686-2.

Z. Yang. Maximum likelihood phylogenetic estimation from DNA sequences with variable rates over sites: approximate methods. Journal of Molecular Evolution, 39(3):306–314, 1994. doi: 10.1007/BF00160154.

C.-H. Yeang, J. F. Darot, H. F. Noller, and D. Haussler. Detecting the coevolution of biosequences—an example of RNA interaction prediction. Molecular Biology and Evolution, 24(9): 2119–2131, 2007.

